# High-frequency phase-switching of *modB* methylase is associated with phenotypic ceftriaxone susceptibility in *Neisseria gonorrhoeae*

**DOI:** 10.1101/2020.04.13.040246

**Authors:** Ola B Brynildsrud, Magnus N Osnes, Kevin C Ma, Yonatan H Grad, Michael Koomey, Dominique A Caugant, Vegard Eldholm

## Abstract

The gonococcal adenine methylases *modA* and *modB*, belonging to separate Type III restriction modification systems, are phase variable and could thus enable rapid adaptation to changing environments. However, the frequency of phase variation across transmission chains and the phenotypic impact of phase variation are largely unknown.

Here we show that the repeat tracts enabling phase variation expand and contract at high rates in both *modA* and *modB*. For *modB*, multiple ON/OFF transition events were identified over the course of a single outbreak.

A mixed effects model using population samples from Norway and a global meta-analysis collection indicates that *modB* in the OFF state is predictive of moderately decreased ceftriaxone susceptibility. Our findings suggest that *modB* orchestration of genome-wide 6-methyladenine modification controls the expression of genes modulating ceftriaxone susceptibility.

**Importance:** Despite significant progress, our current understanding of the genetic basis of antibiotic susceptibility remains incomplete. The gonococcal methylase *modB* is phase variable, meaning that it can be switched ON or OFF via contraction or expansion of a repeat tract in the gene during replication. We find that transitions between the ON and OFF state occur at high frequency. Furthermore, isolates harbouring *modB* in a configuration predicted to be inactive had decreased susceptibility to ceftriaxone, an antibiotic used to treat gonorrhea. This finding improves understanding of the genetic underpinnings of antibiotic resistance, but further work is needed to elucidate the mechanics and broader phenotypic effects of epigenetic modifications and transcription.

## Introduction

*Neisseria gonorrhoeae* has one of the largest known repertoires of restriction modification systems (RMS), with 13-15 systems present in individual genomes (1). The phase variable methylase *modA* has been shown to regulate the expression of genes involved in antigenicity, infection and antibiotic resistance in both *Neisseria meningitidis* and *N. gonorrhoeae* (2). In a study of *N. meningitidis*, locking the methylase *modA11* in ON configuration resulted in significantly higher susceptibility to ciprofloxacin and third generation cephalosporins compared to a strain with *modA11* knocked out (3).

In *N. gonorrhoeae*, two adenine methylases, *modA* and *modB*, which belongs to separate Type III RMS methylases (NgoAXII and NgoAX, respectively), contain genic repeat-tracts resulting in phase variation (2). In addition, the specificity unit *hsdS* of the Type I RMS NgoAV is also phase variable, but phase variation of *hsdS* alters the target motifs that are methylated rather than turning the system ON or OFF (1). Beyond their involvement in restriction modification, these RM systems potentially provide the means for rapid re-arrangements of genome-wide adenine methylation (6mA) patterns and transcription. Here we investigate the frequency of phase variation in *modA* and *modB* and the effect of phase variation on drug susceptibility.

## Results

### *Rapid phase variation of* modB *at a microevolutionary scale*

We investigated the expansion and contraction of phase variable tracts in the Type III RMS methylases *modA* and *modB*, in addition to the Type I RMS specificity unit *hsdS*. First, we selected 10 isolates belonging to the multilocus sequence type (ST) 9363 for nanopore sequencing, as genome-wide methylation load can be read directly from the raw sequencing data (4). For each isolate, hybrid assemblies were generated from Illumina and Oxford Nanopore data using Unicycler (5). Closed genome assemblies were generated for each, ensuring essentially error-free assembly of the RMS modules of interest. This also enabled the direct assessment of target-motif methylation load as a function of RMS status across the 10 isolates.

We found *hsdS* to be invariable in our isolates, suggesting that phase variation of the gene occurs at a relatively modest rate. In contrast, the methylases *modA* and *modB* varied in phase across the 10 ST-9363 isolates. For *modA*, we did not observe a clear effect of predicted ON/OFF status on the methylation load (not shown) on either of the two suggested target motifs identified previously, namely GCAG*A* (1) or AGAAA (2). The gene has a number of putative start codons, which suggests that it might retain some function via other combinations of start codons and tract lengths, even though transcription from alternative start codons upstream of the repeat tract has been reported to be limited (2).

In *modB*, phase variation is mediated by the contraction or expansion of a (CCCAA)n repeat tract (2). There was a clear association between the inferred ON/OFF status and corresponding methylation load on the CC*A*CC target motif (Fig. 1 a). To get a better understanding of the rate of phase variation, we determined the ON/OFF status of the gene in a larger collection of ST-9363 isolates, and mapped the status on a time-resolved phylogeny (see methods). This mapping revealed that *modB* underwent switching between the ON and OFF state at a high rate across the phylogeny, including within a single outbreak spanning less than 3.5 years of evolution (Fig. 1 b).

**Fig. 1.**
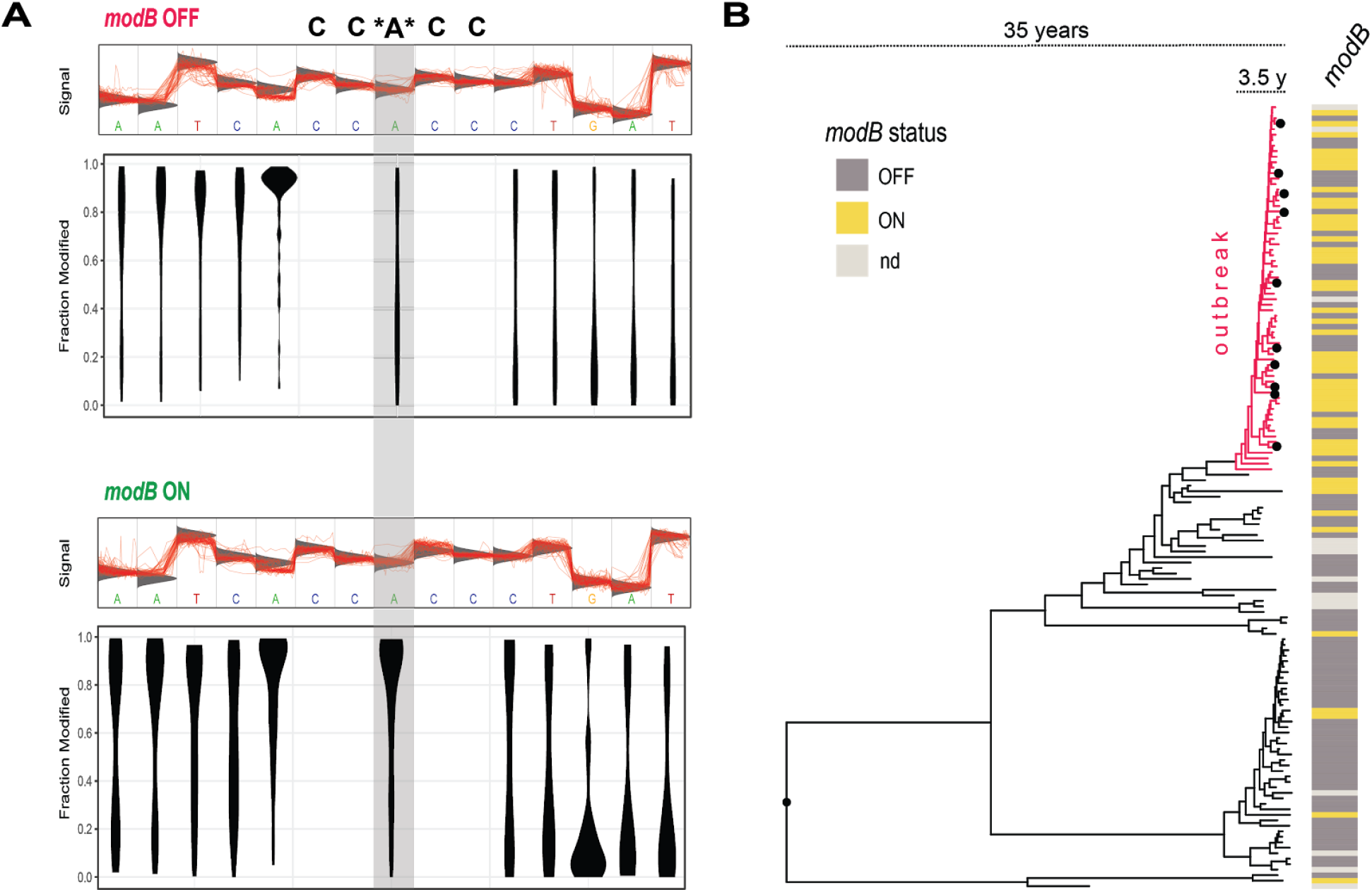
Phase variation of *modB* during the course of an outbreak affects methylation status of CC*A*CC motif. A) Methylation-load on CC*A*CC target sites in isolates stratified by inferred *modB* status (ON/OFF). A “top-heavy” dumbell indicates that a high fraction of the target nucleotides is methylated. B) Time-resolved ST-9363 phylogeny constructed from genome-wide SNPs, annotated with *modB* ON/OFF status. A dense outbreak representing less than 3.5 years of evolution is highlighted in red. Isolates selected for methylome characterization are marked with black dots. As the assemblies were based on Illumina short reads, the repeat tract of the *modB* gene was split between contigs in some assemblies, and the ON/OFF status could thus not be assessed (light grey shading).

### ModB modulates ceftriaxone susceptibility

Next, we investigated whether phase variable methylases could play a role in modulating drug susceptibility. The analyses focused on drugs particularly important for gonorrhea treatment, namely azithromycin and extended-spectrum cephalosporins (cefixime and ceftriaxone). Despite the uncertainties of predicted ModA activity, the encoding gene was included in these analyses.

From a database containing genomes sequenced at the Norwegian Institute of Public Health between 2016 and 2018, we identified a total of 911 isolates with available ceftriaxone and azithromycin minimum inhibitory concentrations (MICs) for which *modB* tract lengths could be determined. To separate methylase effects from known susceptibility determinants segregating in the genetic background, we used linear mixed effect models (see methods).

Even when accounting for known resistance determinants and population structure, *modB* OFF was found to be predictive of a significant, yet modest decrease in susceptibility to ceftriaxone (Fig. 2A, (*modB* ON predictive of a MIC 0.89 (2^-0.17^) times that of *modB* OFF, *i.e*. 11% lower, p = 0.015). No effect was found on cefixime or azithromycin MICs. For *modA*, no significant effects were identified. In line with our finding that *modB* switches between the ON and OFF states at a high rate within ST-9363, the segregation of ON and OFF alleles were not heavily influenced by population structure (□2 p = 0.29 with PopPunk clusters treated as predictor and ON/OFF status as outcome. See Fig. 2B).

**Fig. 2.**
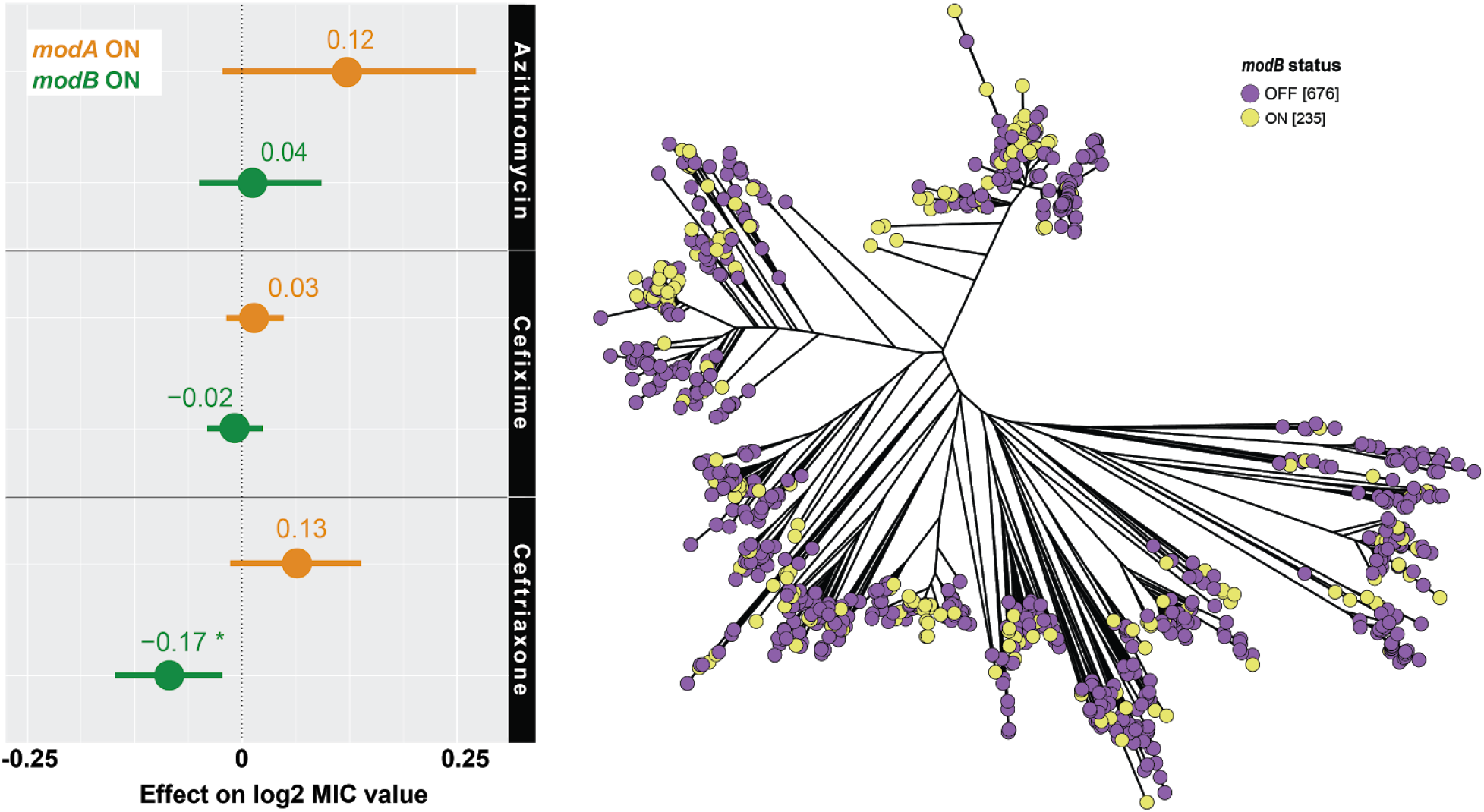
Effect of methylase activity on antibiotic susceptibility. Left panel: Effects of predicted *modA* and *modB* status on antibiotic MIC levels (Point estimate with bars indicating 95% confidence intervals). Right panel: Neighbor joining tree built from core genome pairwise-distances of all isolates with *modB* status mapped to the tips. Due to the uncertainty surrounding the *modA* predictions (see text), the gene status was not mapped on the phylogeny.

To further support the identified effect of ModB on ceftriaxone susceptibility, *modB* ON/OFF status was predicted across a global dataset of 1857 *N. gonorrhoeae* genomes with available MIC data (6). A mixed effect model confirmed a significant effect of *modB* status on ceftriaxone MICs in the verification dataset (*modB* ON effect: −0.11, p = 0.024, corresponding to a 7% reduction in MIC).

## Conclusions and further work

We found that *modB* undergoes rapid phase variation in *N. gonorrhoeae*, with transitions between the ON and OFF states occurring multiple times within a single outbreak. Harboring *modB* in the OFF configuration was predictive of decreased ceftriaxone susceptibility. The effect was modest but significant across two independent genome datasets. No effect was found for *modA*, but these analyses should be repeated in light of more certain methylase activity predictions.

Transcriptomic analyses is needed to define the regulon controlled by *modB* methylation, and functional genetics should be performed to directly assess and validate the role of the *modB* and its regulon in determining drug susceptibility. We note that the ON/OFF status of phase variable genes is invisible to kmer-based genome-wide association studies (GWAS), as most repeat tracts are far longer than typical kmer sizes. Including phase variants in GWAS approaches might improve the sensitivity of these approaches.

## Methods

### Genome analyses

Illumina-only and Illumina + nanopore hybrid assemblies were generated as described previously (7). For analyses of base modifications, nanopore reads were demultiplexed using Deepbinner (8). The methylation load at specific target motifs was quantified from the raw nanopore fast5 reads using Tombo (4).

A ST-9363 whole-genome alignment was generated using parsnp (9), employing a closed hybrid assembly of the ST-9363 isolate 567616 as reference. A whole-genome alignment retaining all reference positions was generated using an in-house script (https://github.com/krepsen/parsnp2fasta). A recombination-corrected maximum likelihood phylogeny was generated using Gubbins (10). The Gubbins output was loaded into BactDating (11). Root-to-tip regression performed in BactDating revealed a sufficient temporal signal (Fig. S1). A time-resolved tree was generated using 4×10^7^ markov chain monte carlo steps (Figs. S2 and S3). The resulting evolutionary rate was 5.67 (95% CI: 3.81-8.28) mutations/genome/year which is similar to earlier studies of *N. gonorrhoeae* (7, 12).

A neighbor-joining phylogeny of all Norwegian *N. gonorrhoeae* genomes (2016-2018) with available phenotypic drug susceptibility data and for which *modB* tract lengths could be determined, was generated using PopPunk and RapidNJ (13, 14). The tree was visualized and edited in GrapeTree (15).

To determine the length of phase variable tracts in *modB* and *hsdS*, these were aligned against the corresponding genes annotated in the FA1090 reference genome using BLAST (16). As we struggled to predict *modA* activity, we chose to use the predictions obtained by aligning assemblies to the *modA* allele-database in pubMLST (17), and only retaining perfect matches for statistical analyses. However, it should be kept in mind that our methylation analyses suggest that further work is needed to determine the actual methylase activity of various *modA* configurations.

A total of 911 genomes for which *modB* state could be determined and for which phenotypic MIC data were available, were included in the Norwegian dataset (supplementary dataset 1).

To account for known resistance determinants, we annotated each genome with information on the following: *mtrD* mosaic (yes/no), *modA* (ON/OFF), modB (ON/OFF), *ponA* (S421L yes/no), *mtrR* (−35A del, A39T and G45D: present/absent), *penA* (pubMLST allele), *porB* (codons 120 and 121 mutated: none, one or both), *23S* (presence/absence of at least one allele harboring A2059G or C2611T) and *rplD* (pubMLST allele). The assignments were based on manual inspection and curation of alignments (*modB* and *mtrD*) or on BLAST searches against pubMLST gene-specific allele databases.

For the global dataset, a total of 1857 genomes (out of 4852) for which *modB* state could be determined and for which phenotypic ceftriaxone MIC data were available were included (supplementary dataset 2). Resistance determinants were acquired from the previous study (6), which identified the following known resistance determinants: *mtrR* (−35A del, A39T and G45D: present/absent, and loss of function mutations), *porB* (codons 120 and 121 mutated: none, one or both), *penA* (codons 501, 542, 551, and BAPS allele classifications), *ponA* (codon 421), and *mtrC* (loss of function mutation).

### Mixed effect models

The linear mixed effect models built for the Norwegian dataset were set up as follows: For the ceftriaxone model, *modA, modB, mtrD, ponA, mtrR* and *porB* assignments were treated as fixed effects, whereas *penA* alleles were included as a random effect due to the high number of alleles. For the azithromycin model, *modA, modB, mtrD* and *mtrR* assignments were included as fixed effects, whereas *rplD* allele assignments were treated as a random effect based on the same reasoning. In addition, genomes were clustered with Poppunk (13), and cluster assignments used as a random effect in both models to take into account any unknown factors associated with population structure. The outcome variables, ceftriaxone and azithromycin MICs, were log_2_-transformed. For fixed effect variables, the wildtype allele was treated as the reference category. Variable inclusion was determined in a forward stepwise process, and nested models were evaluated by the Bayesian information criterion (BIC). P-values were estimated using the lme4 package (https://cran.r-project.org/web/packages/lme4/index.html). The full output of estimated beta-values for the fixed effects are available in Figures S4-S6.

The linear mixed effect model employed on the global verification dataset was set up similarly, as follows: *modB* state, *mtrR* alleles, *ponA* alleles, *penA* alleles and *porB* alleles were treated as fixed effects, whereas country of isolation, *penA* BAPS cluster (for identifying mosaic alleles), and overall BAPS cluster (to account for overall population structure) were treated as random effects. The lmerTest package (https://cran.r-project.org/web/packages/lmerTest/index.html) was used as an approximate method to estimate p-values.2

## Supporting information

supplementary_dataset1_NIPH

supplementary_dataset2_Harvard

## Acknowledgements

The authors would like to thank our colleagues responsible for culturing and whole genome sequencing of gonococci at the Norwegian Institute of Public Health. Magnus N. Osnes is supported by The National Graduate School in Infection Biology and Antimicrobials (IBA) hosted by the University of Oslo. YHG is supported by NIH/NIAID grant R01 AI132606. KCM is supported by the NSF GRFP.

This publication made use of the Neisseria Multi Locus Sequence Typing website (https://pubmlst.org/neisseria/) sited at the University of Oxford and funded by the Wellcome Trust and European Union.

## Notes

### Competing Interest Statement

The authors have declared no competing interest.

### Summary of Updates

In this revised version, we used mixed effect models to assess the effect of modA and modB status on phenotypic antibiotic MICs. We also included a second global verification dataset to confirm out findings from the Norwegian dataset.

